# The Extracellular Matrix Regulates Invasion in Fusion-Negative Rhabdomyosarcoma via YAP-PIEZO1 Signaling Axis

**DOI:** 10.64898/2026.01.05.697722

**Authors:** Yuanzhong Pan, Juha Kim, Brian M. Wong, Esteban Espuny, JinSeok Park

## Abstract

**Background:** Fusion-negative rhabdomyosarcoma (FNRMS) represents the most prevalent subtype of rhabdomyosarcoma, the most common pediatric soft tissue sarcoma. Although its invasion is a leading cause of recurrence and poor prognosis, its underlying mechanism remains unclear. We investigated how extracellular matrix (ECM) density regulates FNRMS progression via mechano-transduction.

**Methods:** We used three-dimensional spheroid invasion assays with RD and SMS-CTR cells embedded in varying collagen concentrations. Mechanistic insights were gained through immunofluorescence, ChIP-seq re-analysis, Fluo-4 calcium live cell imaging, and pharmacological inhibition of the YAP–PIEZO1 axis.

**Results:** High ECM density significantly enhanced invasive spreading, correlating with increased YAP nuclear localization. YAP overexpression was sufficient to promote invasion, while its inhibition attenuated the matrix-enhanced phenotype. We identified *PIEZO1* as a direct transcriptional target of the YAP. High ECM density stimulated PIEZO1-dependent calcium influx, which was required for invasion. Furthermore, elevated *PIEZO1* expression was significantly associated with poorer overall survival in FNRMS patients. Targeting YAP or ROCK signaling effectively suppressed both calcium flux and invasion.

**Conclusions:** Our findings establish a YAP–PIEZO1 axis linking ECM density to FNRMS invasion. This mechanosensitive pathway represents a potential therapeutic vulnerability in aggressive FNRMS.

**Simple Summary:** Fusion-negative rhabdomyosarcoma (FNRMS) is the prevalent subtype of rhabdomyosarcoma, the most common pediatric soft tissue sarcoma. Its invasion often leads to recurrence and poor prognosis. This study investigates how the density of the extracellular matrix (ECM), surrounding cancer cells in the tissue, influences FNRMS invasion. We found that high ECM density activates the protein YAP, which directly triggers the expression of the mechanical sensor PIEZO1. This sensor allows calcium to enter the cells, providing a signal that facilitates invasion. Pharmacological inhibition of this axis successfully reduced the invasive potential, highlighting a novel therapeutic vulnerability for FNRMS patients.

## Introduction

Rhabdomyosarcoma (RMS) is the most common soft-tissue sarcoma of childhood and adolescence, arising from skeletal muscle progenitor cells that undergo malignant transformation^1^. Despite advances in multimodal therapy, clinical outcomes remain suboptimal, particularly for patients with high-risk disease or recurrent tumors. RMS comprises two major molecular subtypes with distinct clinical behaviors: fusion-positive (FPRMS), characterized by chromosomal translocations such as PAX3–FOXO1 or PAX7– FOXO1, and fusion-negative (FNRMS), which lacks these fusions. Although FPRMS has historically been associated with higher aggressive disease, FNRMS accounts for approximately 60% of RMS cases and remains incompletely understood at the mechanistic level^2^. Consequently, a deeper understanding of the biological underpinnings of FNRMS is essential for improving therapeutic strategies for this clinically challenging subtype.

Tumor progression is governed not only by cell-intrinsic oncogenic programs but also by the biophysical and biochemical properties of the tumor microenvironment (TME) ^3, 4^. Among these, the extracellular matrix (ECM) plays a critical role in regulating cancer cell migration, invasion, and survival^5–7^. Changes in ECM composition, architecture, and stiffness are hallmarks of many solid tumors and can profoundly influence cellular behaviors through mechano-transduction pathways. While increasing evidence suggests that RMS cells respond to the ECM^8–11^, how FNRMS senses and adapts to the ECM remains poorly defined. Given that the invasion is a major clinical concern in RMS, elucidating the mechanobiology of FNRMS may uncover previously unrecognized vulnerabilities.

A key effector of mechanical signaling is the transcriptional co-activator Yes-associated protein (YAP), a central component of the Hippo pathway ^12–15^. YAP shuttles between the cytoplasm and nucleus in response to increased ECM density, where it regulates gene expression programs that control proliferation, survival, and migration. YAP has been implicated in the progression of multiple cancer types. While YAP activity is highly activated in FNRMS and up-regulate pro-proliferative and oncogenic genes, its mechanistic role in FNRMS, particularly in the context of the interaction with the TME, remains unclear. Understanding whether YAP functions as a mediator that bridges environmental ECM cues to FNRMS cell invasive behaviors associated with clinical outcomes could provide critical insight into potential therapeutic intervention points.

In this study, we investigate how ECM density modulates the invasive behavior of FNRMS cells and test the hypothesis that YAP serves as a mechano-transduction regulator of this process. We identify PIEZO1, a mechanosensitive ion channel that facilitates calcium influx ^16–18^, as a direct downstream target of YAP. Our findings demonstrate that the ECM-YAP–PIEZO1 axis regulates calcium signaling to promote FNRMS invasion. Furthermore, we determine that repression of this regulatory axis reduces the invasive potential of FNRMS cells, identifying potential therapeutic targets for FNRMS.

## Results

### FNRMS cells exhibit enhanced invasive behavior on high ECM density

Multiple cancer types exhibit increased ECM density in high-risk or metastatic subgroups compared to low-risk or local counterparts ^19–21^. Furthermore, elevated ECM density has been shown to promote tumor cell invasion ^22, 23^. To determine whether the ECM density similarly influences the migratory behavior of FNRMS cells, we performed a three-dimensional (3D) spheroid invasion assay using collagen type I matrices at varying concentrations. Spheroids of uniform sizes were generated from two FNRMS cell lines, RD and SMS-CTR, and seeded onto collagen gels at low (0.5%) or high (1.0%) concentrations, hereafter referred to as low- and high-density matrices, respectively.

Live-cell imaging revealed that spheroids from both cell lines exhibited greater radial expansion, reflecting the invasion into the surrounding matrix, when cultured on high-density collagen gels compared to low-density conditions (Figure 1A–B). This enhanced outgrowth was consistent across spheroids of different initial sizes, indicating that the effect was independent of initial spheroid size. Quantitative analysis of spheroid spreading area over multiple time points demonstrated a consistently larger invaded area on high-density matrices for all conditions tested (Figure 1C). Collectively, these results indicate that increased collagen matrix density enhances the invasive potential of FNRMS cells in a 3D context.

**Figure 1.**
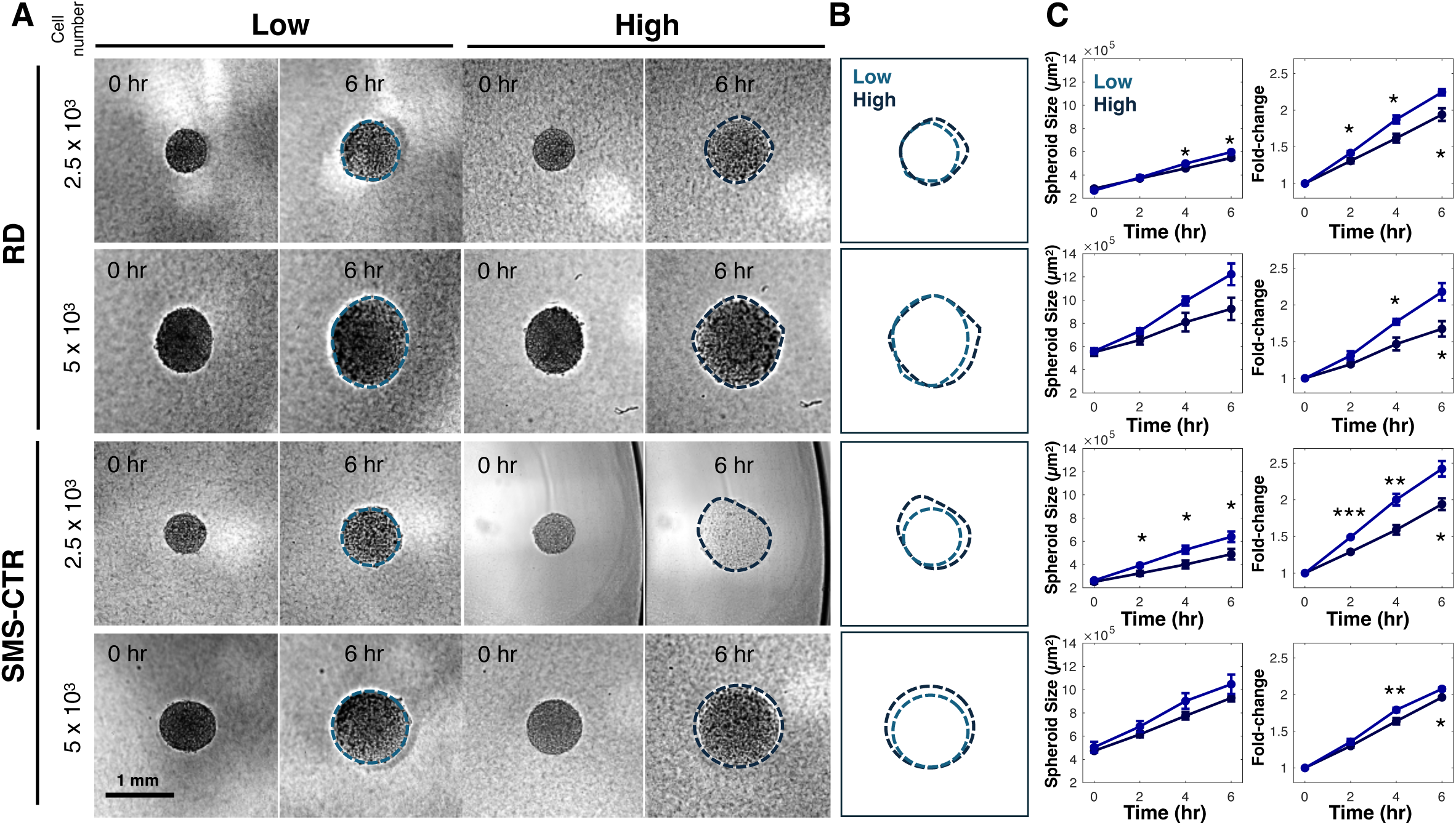
Increased ECM density promotes FNRMS spheroid invasion. **A.** Representative images of FNRMS spheroids. RD and SMS-CTR spheroids of varying initial sizes (2.5 x 10^3^ and 5 x 10^3^ cells per spheroid) were seeded on low-density (0.5% collagen type I) or high-density (1.0% collagen type I) matrices. Images show the same spheroids at 0 and 6 hours (h) post-seeding. Dashed lines indicate the quantified boundaries of the spheroids. Scale bar = 1 mm. **B.** Representative outlines of spheroids to illustrate the extent of radial expansion in low (light blue) versus high (dark blue) density conditions. **C.** Quantitative analysis of invasive outgrowth. (Left) Longitudinal measurement of spheroid area over 6 h. (Right) Normalized fold-change in spheroid area relative to t = 0 h. Error bars represent SEM from two or three independent spheroids. Statistical significance comparing low- and high-density groups at each time point was assessed using two-sided Student’s t-tests: *p < 0.05, **p < 0.01, ***p < 0.001.

### YAP mediates matrix-dependent invasion of FNRMS cells

Cancer cell invasion is regulated by mechano-transduction signaling pathways that enable cells to sense and respond to the physical properties of the ECM ^24–26^. The transcriptional co-activator YAP is a central effector of mechano-transduction and has been implicated in ECM density-dependent regulation of cell invasion in multiple cancer types. We therefore investigated whether YAP mediates the enhanced invasive behavior of FNRMS cells observed on high-density collagen matrices. To assess YAP activity in response to matrix properties, we performed immunofluorescence staining for YAP in FNRMS spheroids cultured on low- or high-density collagen gels.

YAP activity is regulated by its subcellular localization: upon activation, YAP translocates to the nucleus, where it functions as a transcriptional co-activator. Thus, we quantified nuclear YAP signal intensity in cells located near the invasive front of spheroids. Both RD and SMS-CTR spheroids, across all sizes examined, exhibited significantly increased nuclear YAP accumulation when cultured on high-density collagen matrices compared to low-density conditions (Figure 2A–B). These data indicate that increased matrix density elevates YAP activity in invading FNRMS cells.

**Figure 2.**
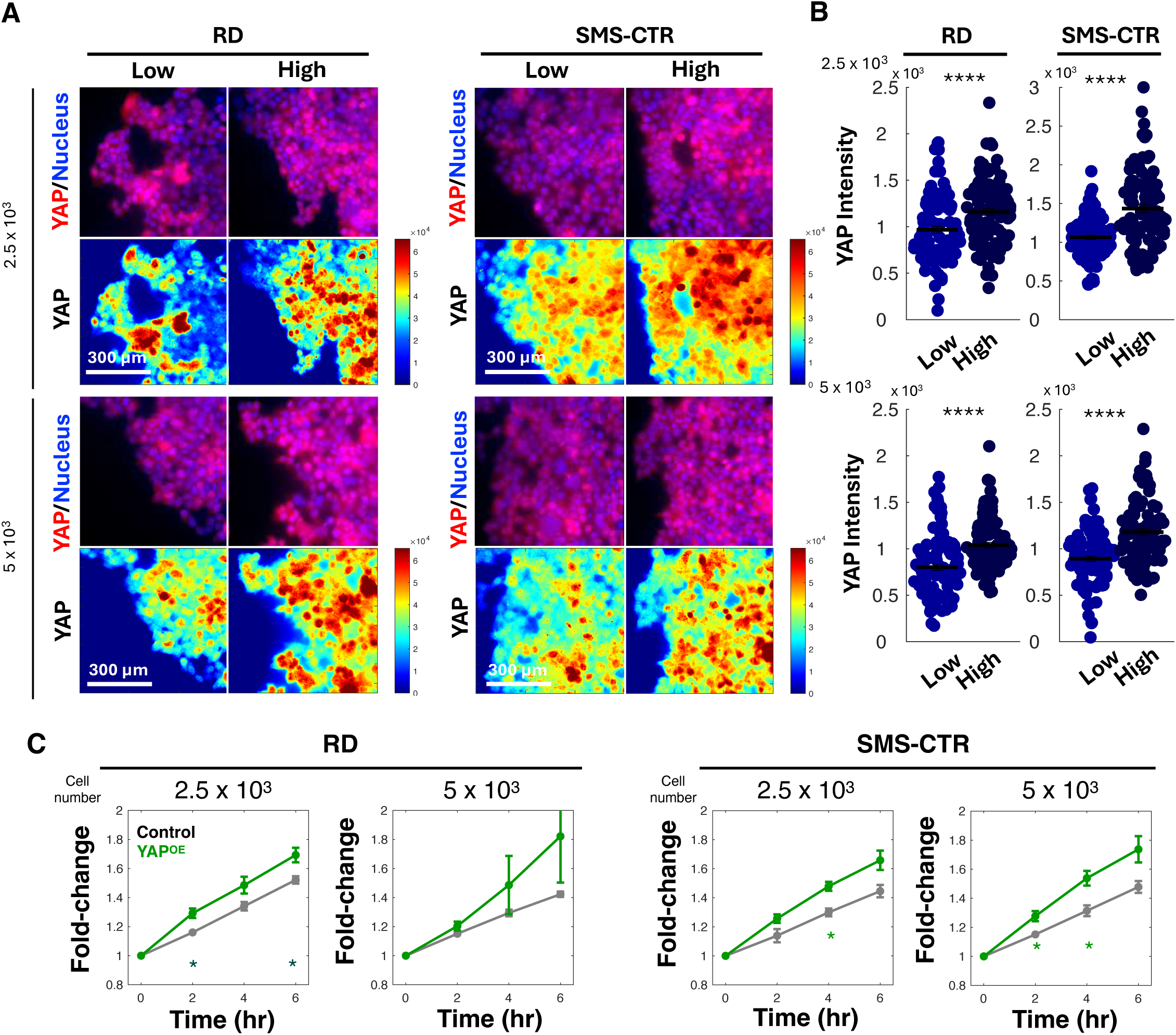
YAP mediates ECM density-dependent invasion in FNRMS cells. **A.** Representative immunofluorescence images of YAP localization in RD and SMS-CTR spheroids (initial sizes: 2.5 x 10^3 and 5 x 10^3 cells) on low- or high-density collagen matrices. Panels show YAP (red) merged with nuclei (DAPI, blue), alongside corresponding pseudo-color heatmaps representing YAP signal intensity. **B.** Quantification of nuclear YAP signal intensity. Data represent the nuclear YAP intensity measured in cells at the invasive front of the spheroids. Horizontal lines indicate the mean value for each group. Error bars represent SEM. ****p < 0.0001 by two-sided Student’s t-test comparing low- and high-density conditions. **C.** Effect of YAP overexpression on FNRMS invasion. Normalized fold-change in spheroid area (relative to t = 0 h) for Control and YAP-OE FNRMS cells over a 6 h period. Normalized fold-change in spheroid area relative to t = 0 h. Error bars represent SEM from two or three independent spheroids. Statistical significance comparing Control and YAP-OE groups at each time point was assessed using two-sided Student’s t-tests: *p < 0.05.

To determine whether YAP activation is sufficient to promote invasion, we generated YAP-overexpressing (YAP-OE) FNRMS cells and assembled spheroids for invasion assays. Quantitative analysis of spheroid spreading area over time revealed that YAP overexpression significantly increased the invasive spreading relative to control spheroids (Figure 2C). This effect phenocopied the enhanced invasion observed on high-density collagen matrices, supporting a functional role for YAP in driving ECM-dependent invasion. Together, these results demonstrate that YAP activity is elevated in response to increased matrix density and that YAP is sufficient to promote invasive behavior in FNRMS spheroids, identifying YAP as a key mediator of stiffness-dependent invasion in this disease context.

### PIEZO1 functions as a downstream effector of YAP signaling in FNRMS cells

Because YAP primarily functions as a transcriptional co-activator, we next sought to identify downstream effectors through which YAP regulates ECM density-dependent invasion in FNRMS. In particular, we focused on genes involved in mechano-transduction that could link ECM density to intracellular signaling programs.

PIEZO1 is a mechanosensitive calcium-permeable ion channel that transduces mechanical cues from the extracellular environment into intracellular calcium signaling^16–18^. Given that calcium signaling promotes cancer cell invasion^27–30^, PIEZO1-mediate calcium signaling has been implicated in cancer cell invasion. Tremblay et al. demonstrated that YAP activation drives oncogenic transformation of satellite cells, which exhibit hallmark features of FNRMS ^31^. To determine whether PIEZO1 is a transcriptional target of YAP-TEAD1(the primary DNA binding partner of YAP ^31^) in FNRMS, we reanalyzed publicly available datasets of chromatin immunoprecipitation sequencing (ChIP-seq) for TEAD1 of mouse satellite cells transformed via YAP activation and RD cells, both of which serve as robust models for FNRMS. Our analysis revealed prominent YAP-TEAD1 occupancy within the proximal promoter region of the *PIEZO1* gene in both mouse and human FNRMS cells (Figure 3A). These findings suggest that YAP directly regulates *PIEZO1* transcription in FNRMS.

**Figure 3.**
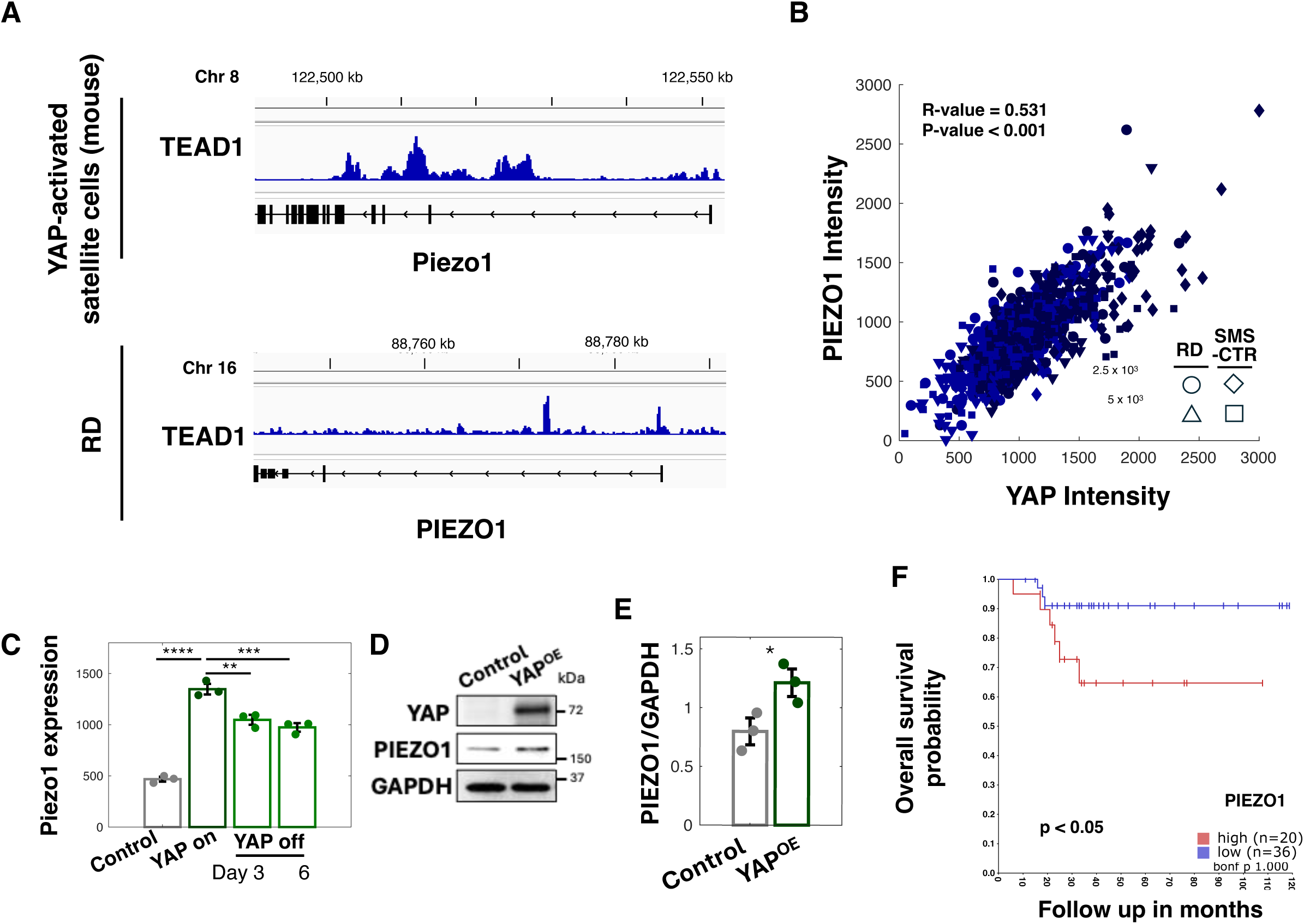
PIEZO1 is a direct downstream target of YAP signaling in FNRMS. **A.** ChIP-seq data shows TEAD1 occupancy at the proximal promoter region of the *PIEZO1* gene in mouse satellite cells transformed by YAP activation (top) and human RD FNRMS cells (bottom). Data from GSE55186. **B.** Quantitative analysis of single-cell immunofluorescence data showing a positive linear correlation between nuclear YAP intensity and PIEZO1 protein expression in RD and SMS-CTR cells from co-immunofluorescence staining. The correlation coefficient (R-value) and p-value are calculated using Pearson’s correlation coefficient. **C.** *PIEZO1* transcript levels in FNRMS cells expressing a doxycycline-inducible constitutively active YAP mutant (YAP-ON). Expression was measured in control (non-induced), induced (YAP-ON), and withdrawal (YAP-OFF) conditions at days 3 and 6. Error bars are SEM (n = 3). ****p < 0.0001, ***p < 0.001, **p < 0.01 by two-sided Student’s t-test. **D.** YAP and PIEZO1 protein levels in control and YAP-overexpressing (YAP-OE) SMS-CTR cells. GAPDH serves as a loading control. **E.** Quantification of PIEZO1 protein abundance normalized to GAPDH. Error bars are SEM (n = 3). *p < 0.05 by two-sided Student’s t-test. **F.** Kaplan-Meier overall survival analysis of FNRMS patients stratified by *PIEZO1* expression level (high vs. low). Data were analyzed using the R2 Genomics Analysis and Visualization Platform. The number of patients in each group and the Log-rank test p-value are indicated.

Consistent with this genomic observation, quantitative analysis of immunofluorescence imaging data demonstrated a strong positive linear correlation between nuclear YAP intensity and PIEZO1 expression at the single-cell level (Figure 3B). To further determine whether YAP activity regulates *PIEZO1* expression, we analyzed RNA-seq data from FNRMS cells using a doxycycline-inducible system to express a constitutively active YAP mutant (YAP-ON). Induction of YAP signaling resulted in a significant upregulation of *PIEZO1* transcript levels compared to non-induced control cells (Figure 3C). Notably, this upregulation was reversible, as *PIEZO1* expression returned to baseline levels following the withdrawal of the induction (YAP-OFF).

We next validated these findings at the protein level. Western blot analysis revealed elevated PIEZO1 expression in YAP-OE FNRMS cells relative to controls (Figure 3D). Quantification across multiple independent experiments confirmed a statistically significant increase in PIEZO1 abundance upon YAP overexpression (Figure 3E). Together, these data establish that *PIEZO1* is a downstream target of YAP-mediated transcriptional regulation in FNRMS cells.

Finally, analysis of clinical FNRMS patient datasets revealed that elevated *PIEZO1* expression is significantly associated with poorer overall survival (Figure 3F), underscoring the clinical relevance of PIEZO1 in FNRMS progression. These findings suggest that the ECM-YAP–PIEZO1 axis contributes to the invasive and aggressive behavior of FNRMS tumors and warrants further investigation.

### PIEZO1-dependent calcium signaling promotes invasion of FNRMS cells on high-density matrices

To determine whether PIEZO1 functionally contributes to ECM density-dependent invasion, we first examined its expression in invading FNRMS spheroids. Immunofluorescence co-staining of PIEZO1 along with YAP revealed significantly elevated PIEZO1 signal intensity in cells within spheroids cultured on high-density collagen I matrices compared to low-density conditions (Figure 4A–B), consistent with increased YAP activity under these conditions. Because PIEZO1 functions as a mechanosensitive, calcium-permeable ion channel, we next assessed whether ECM density modulates intracellular calcium signaling. Live-cell calcium flux imaging using the Fluo-4 AM dye was performed on FNRMS spheroids cultured on low- or high-density collagen I gels (Figure 4C). Quantitative analysis demonstrated a significantly higher calcium influx in spheroids cultured on high-density matrices (Figure 4D), indicating that increased matrix concentration enhances calcium signaling in FNRMS cells.

**Figure 4.**
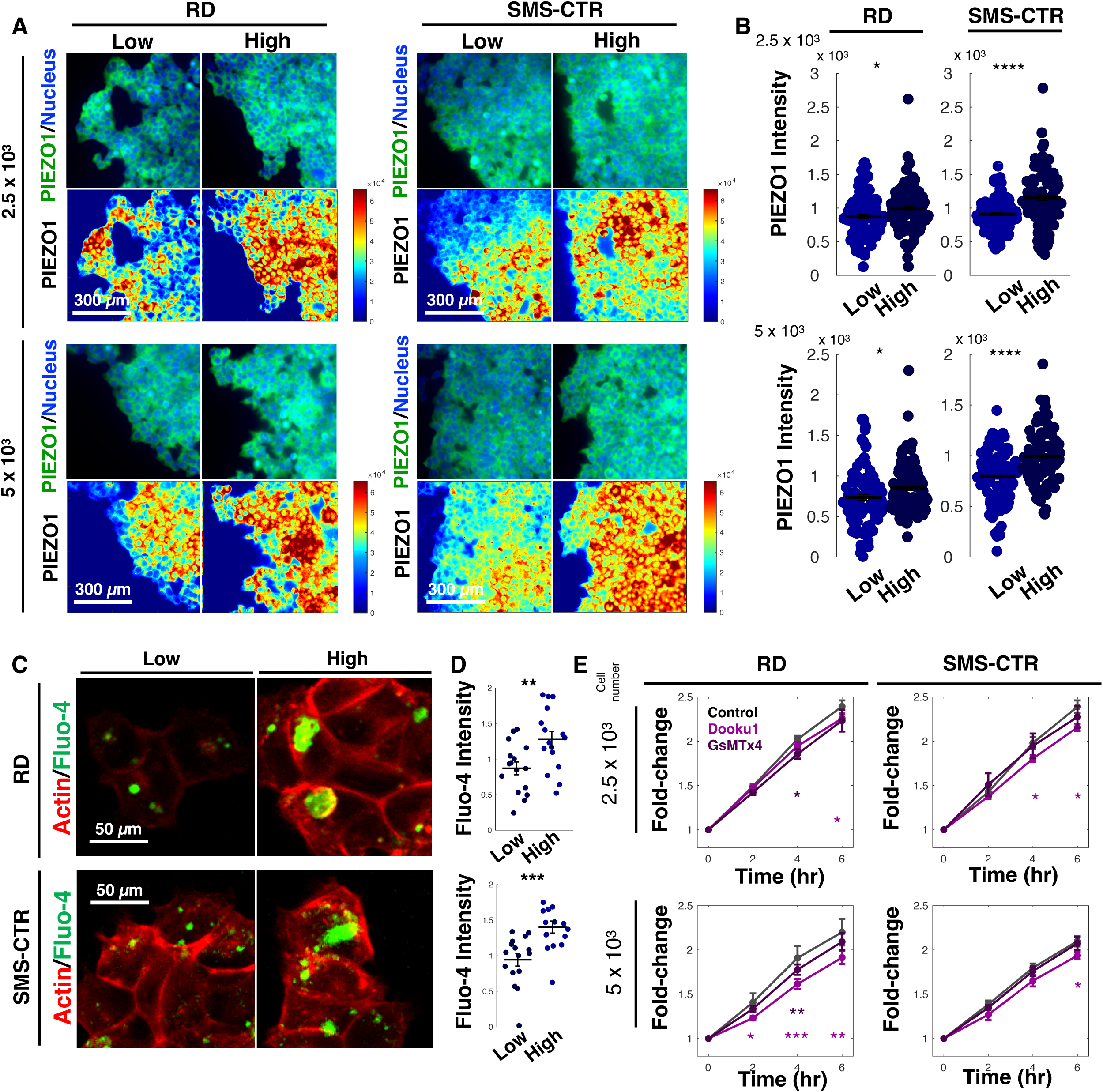
PIEZO1-mediated calcium signaling drives ECM density-dependent invasion in FNRMS. **A.** Representative immunofluorescence images of PIEZO1 localization in RD and SMS-CTR spheroids cultured within low- or high-density collagen matrices. Top panels display PIEZO1 (green) merged with nuclei (DAPI, blue). Bottom panels show corresponding pseudo-color heatmaps representing PIEZO1 signal intensity. **B.** Quantification of PIEZO1 signal intensity from the images in Panel A. Data points represent individual cells measured at the invasive front. Horizontal lines indicate the mean value for each group. ****p < 0.0001, *p < 0.05 by two-sided Student’s t-test. **C.** Representative live-cell calcium imaging of RD and SMS-CTR cells embedded in low- or high-density collagen gels. Cells were loaded with the calcium indicator Fluo-4 AM (green) and stained for F-actin (red). Increased green fluorescence indicates elevated intracellular calcium levels. **D.** Quantification of intracellular calcium influx (Fluo-4 intensity) from Panel C. Data points represent individual cells. Horizontal lines indicate the mean. ***p < 0.001, **p < 0.01 by two-sided Student’s t-test. **E.** Quantitative analysis of spheroid invasion over 6 hours in RD and SMS-CTR spheroids treated with vehicle control, Dooku1 (5 μM), or GsMTx4 (5 μM). Data show the normalized fold-change in spheroid area relative to t = 0 h. Error bars represent SEM from two or three independent spheroids. Statistical significances compared to the control group at the indicated time points were evaluated using two-sided Student’s t-test: *p < 0.05, **p < 0.01

To evaluate the functional contribution of PIEZO1-mediated calcium signaling to spheroid invasion, we pharmacologically inhibited PIEZO1 by the treatment of Dooku1 (PIEZO1 antagonist) ^32^ and GsMTx4 (a mechanical channel blocker) ^33^. PIEZO1 inhibition resulted in in a significant reduction in spheroid spreading area compared to vehicle-treated controls (Figure 4E). Collectively, these data indicate that increased ECM density correlates with elevated PIEZO1 expression and enhanced calcium signaling, which in turn facilitates the invasive behavior of FNRMS cells. These findings support a model in which PIEZO1-mediated calcium signaling acts as a key effector downstream of YAP to drive ECM density-dependent invasion.

### Pharmacological inhibition of YAP signaling attenuates calcium influx and spheroid

To evaluate the potential therapeutic relevance of the mechano-transduction pathways identified in this study—specifically the YAP–PIEZO1 signaling axis—we tested whether pharmacological inhibition of YAP-associated signaling could suppress the elevated calcium influx and invasive behavior observed in FNRMS spheroids cultured in high-density collagen matrices. Verteporfin has been utilized to disrupt the interaction between YAP and TEAD transcription factors, thereby inhibiting YAP-dependent transcriptional activity^34^. Additionally, we employed Y-27632, a potent inhibitor of Rho-associated protein kinase (ROCK), which reduces YAP nuclear localization by modulating cytoskeletal tension in response to the ECM density^35, 36^.

Live-cell imaging revealed that treatment with either Verteporfin or Y-27632 significantly reduced intracellular calcium influx in FNRMS cells compared to vehicle-treated controls (Figure 5A–B). Consistent with these results, quantitative analysis of spheroid invasion assays demonstrated that both inhibitors significantly attenuated the spheroid spreading area over time (Figure 5C). While invasion was not completely abolished, pharmacological inhibition of the YAP signaling axis markedly suppressed the ECM density-enhanced invasive phenotype. Collectively, these results demonstrate that targeting the YAP–PIEZO1– calcium axis can effectively reduce the invasive capacity of FNRMS cells, highlighting its potential as a therapeutic vulnerability in the context of high ECM density.

**Figure 5.**
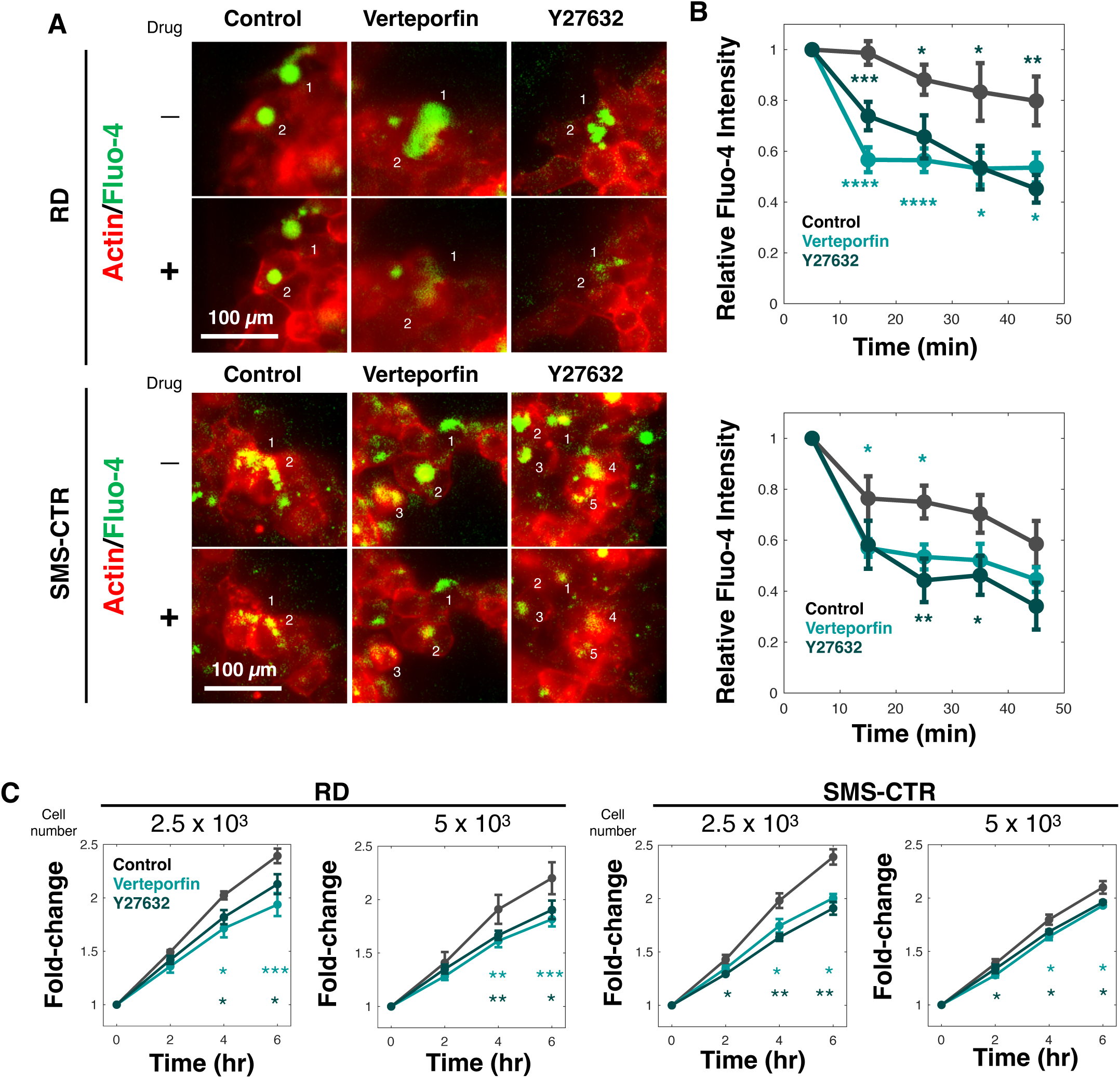
Targeting YAP signaling suppresses matrix-dependent calcium signaling and invasion. **A.** Representative live-cell calcium imaging of RD and SMS-CTR cells embedded in high-density collagen gels. Cells were treated with DMSO (vehicle control), the YAP inhibitor Verteporfin, or the ROCK inhibitor Y-27632. Cells were stained with the calcium indicator Fluo-4 AM (green) and F-actin (red). Decreased green fluorescence indicates reduced intracellular calcium levels in inhibitor-treated cells. **B.** Quantification of intracellular calcium (Fluo-4 intensity) from the experiments represented in Panel A. Error bars are SEM. Ten cells per group were analyzed: ***p < 0.001 compared to the DMSO control group by two-sided Student’s t-test. **C.** Quantitative analysis of spheroid invasion over 6 hours in high-density matrices. RD and SMS-CTR spheroids were treated with DMSO Control, Verteporfin, or Y-27632. Data show the normalized fold-change in spheroid area relative to t = 0 h. Error bars represent SEM from independent two or three spheroids. *p < 0.05, **p < 0.01 compared to the control group at the indicated time points by two-sided Student’s t-test.

## Discussion

In this study, we demonstrate that ECM density enhances the invasive behavior of FNRMS cells and identify a mechano-transduction pathway linking matrix density to tumor cell invasion, ECM-YAP-PIEZO1 signaling axis. Using 3D spheroid invasion models, we show that high-density collagen matrices promote increased spheroid spreading, indicating elevated invasive potential. These findings highlight the critical role of ECM properties in the ECM in regulating FNRMS cell behavior and underscore the necessity of considering microenvironmental cues to fully understand disease progression.

Mechanistically, we identify YAP as a key mediator of ECM density-dependent invasion in FNRMS. Increased matrix density was associated with elevated nuclear YAP localization in invading cells, and YAP overexpression was sufficient to enhance spheroid invasion. These observations are consistent with YAP’s established role as a central effector of mechano-transduction pathway and extend its functional relevance to FNRMS, a molecular subtype that is more prevalent than fusion-positive disease yet remains mechanistically understudied. Our data suggest that YAP integrates mechanical signals from the ECM to drive transcriptional programs that promote invasive behavior in this rhabdomyosarcoma subtype.

We further identify the mechanosensitive calcium channel PIEZO1 as a downstream effector of YAP signaling in FNRMS cells. ChIP-seq reanalysis, transcriptomic profiling, and protein-level validation collectively establish *PIEZO1* as a YAP-regulated gene. Elevated *PIEZO1* expression correlated with increased YAP activity and was significantly associated with poorer clinical outcomes in FNRMS patients, underscoring potential relevance in FNRMS. Functional analyses demonstrated that high-density matrices increase PIEZO1 expression and calcium influx in spheroids, directly linking mechanical cues to intracellular calcium signaling. Furthermore, pharmacological inhibition of PIEZO1 cause significant reductions in invasion. These data indicate that PIEZO1 positively contributes to ECM density -dependent invasive behavior, consistent with the involvement of parallel or compensatory mechano-transduction pathways.

Importantly, pharmacological inhibition of YAP-associated signaling attenuated both calcium influx and spheroid invasion, supporting the central role of YAP in coordinating mechanical signaling outputs in FNRMS. Although the inhibitors used in this study are not clinically optimized for sarcoma treatment, these experiments provide proof-of-concept that targeting YAP-centered mechano-transduction pathways can suppress FNRMS invasion.

Together, our findings support a model in which ECM density promotes YAP activation, leading to transcriptional upregulation of mechanosensitive effector PIEZO1 and enhanced calcium signaling, ultimately facilitating FNRMS cell invasion. Overall, this work reveals a previously unappreciated YAP–PIEZO1 signaling axis that links ECM mechanics to invasive behavior in FNRMS. These findings advance our understanding of FNRMS mechanobiology and suggest that targeting mechano-transduction pathways may represent a complementary therapeutic strategy to limit invasion in this aggressive pediatric cancer.

## Materials and Methods

### Cell Culture

FNRMS cell lines SMS-CTR and RD were obtained from Childhood Cancer Repository. Cells were cultured in Dulbecco’s Modified Eagle Medium (DMEM, Gibco cat. 11965-092) supplemented with 20% Fetal Bovine Serum (FBS, Gibco cat. A5670701) and 1% penicillin/streptomycin (Gibco cat. 15140-122), in a 37 °C, 5% CO_2_ incubator.

### Spheroid Formation and Collagen Invasion Assay

To generate spheroids, the liquid overlay method was employed. Adherent cell cultures were detached using Accutase (Millipore cat. SCR005), followed by dilution with culture media to allow for the seeding of 5,000 and 2,500 cells per 100 µL cell culture media, which were deposited in a 96-well spheroid plate (Corning cat. 4515). The cells were maintained in a 37°C, 5% CO_2_ incubator for 24 hours until compact aggregated spheroids were formed.

For collagen invasion assay, a stock solution of type I collagen from rat tail (Gibco cat. A10483-01) was used to prepare collagen gel at a final concentration of 0.5% and 1.0%. In a sterile tube, for each 400 µL of final gel solution, 253 µL (187 µL for 1.0% gel) of H_2_O, 40 µL of HEPES (Sigma cat. H0887), 40 µL of DMEM, 6.8 µL of NaOH (Thermo Scientific cat. 047124.K2), and 67 µL (134 µL for 1% gel) of collagen stock solution were combined to make the gel mixture, and 40 µL of the mixture was deposited into each well of a 96-well plate or an 8-chamber cover slide (Thermo Scientific cat. 155409). The plate was incubated at 37°C for 20 min to allow the gel to solidify. The spheroids were then collected using a standard pipette tip with 3 mm of the end cut off and deposited onto the collagen gel.

Live-cell imaging was conducted to visualize the invasion of spheroids on a Nikon Eclipse C2 inverted microscopy that is equipped with an OKO atmosphere controlling system to maintains humidity, 37°C temperature, and 5% CO_2_. The spreading area was quantified using ImageJ.

### Immunofluorescent (IF) Staining and Imaging

For IF staining, media were removed from cultured spheroids or cell monolayer, which was washed with PBS twice. Cells were fixed with 4% formaldehyde at room temperature for 20 mins, then washed twice. PBS containing 0.1% Triton X-100 (Thermo Scientific cat. J66624-AE) was added to permeabilize the membrane for 45 mins at room temperature. Blocking was performed with PBS containing 3% BSA (Sigma cat. A9647) for 45 mins, followed by primary antibody (YAP Rabbit antibody, CST cat. 14074; PIEZO1 mouse antibody, Invitrogen cat. MA5-32876; 1:200 dilution) staining for 1 hour at room temperature or overnight at 4°C. After two washes, secondary antibodies (AF594 goat anti-rabbit IgG, Invitrogen cat. A-11012; AF488 goat anti-mouse IgG, Invitrogen cat. A-10680; 1:1,000 dilution) and nucleus dye (Hoechst 33342, Invitrogen cat. 62249, 1:10,000 dilution) diluted in PBS containing 1% BSA was added and incubated at room temperature in dark for 45 mins. The cells were washed three times with PBS and imaged with a Nikon fluorescent microscope. Quantification of IF intensities was performed using custom-made MATLAB algorithm (MathWorks) and ImageJ.

### Sequencing data re-analysis

ChIP-seq data from YAP-activated satellite and RD cells were obtained from GSE55186. Additionally, RNA-seq data from doxycycline-inducible YAP1 S127A-driven FNRMS model tumors and tumors regressed following doxycycline withdrawal were downloaded from GSE47198. Processed data and genomic tracks were visualized using the Integrative Genomics Viewer (IGV).

### YAP Overexpression

YAP overexpression was performed by expressing YAP-expressing plasmid (mEGFP-N1-YAP, addgene cat. 166457) using Lipofectamine 3000 Transfection kit (Invitrogen cat. L3000-015) following to the manufacturer’s protocol. Briefly, for each transfection reaction in one well of a 6-well plate, 7 µL was added to 125 µL OptiMEM (Gibco cat. 51985034) followed by vortexing for 3 seconds. In a separate 1.7 mL tube, 5 µg of plasmid DNA was added to 125 µL OptiMEM, then 10 µL of P3000 to the mixture. The lipofectamine solution is then added to the DNA solution, mixed by pipetting, and incubated at room temperature for 10 mins. 250 µL of the transfection reaction mixture was then added dropwise to cultured cells that are over 90% confluent. For spheroid experiments, cells were harvested 24 hours after transfection and spheroids were made as described in the previous section. For western blot, cells were harvested 36 hours after transfection.

### Western Blotting

For western blot, cells were harvested with Accutase (Millipore cat. SCR005) after 36 hours of transfection, washed twice with PBS, and lysed with RIPA buffer (Thermo Scientific cat. 89901) containing 1X protease and phosphate inhibitor cocktail (Thermo Scientific cat. 78440). Cell lysates were centrifuged at 20,000 g for 15 min at 4°C to pellet insoluble portion of the lysate, only supernatant was kept. Protein concentration was quantified via BCA assay using Pierce BCA kit (Thermo Scientific cat. 23227) and a Varioskan plate reader, and all samples were normalized to the same concentration.

5 µg of protein was loaded on a 15-well gel (BioRad cat. 4561096) and electrophoresis was run in 1X Tris/Glycine/SDS buffer (diluted from 10X stock, BioRad cat. 1610772) at 50 V for 3 hours at room temperature. Semidry transfer was performed on a BioRad Trans-Blot Turbo transfer system at 25 V for 10 min. Membrane was then cut and incubated in 3% BAS in Tris-buffered saline containing 1% Tween-20 (TBST) solution at room temperature for 1 hour to block unspecific binding. Primary antibodies (YAP Rabbit antibody, CST cat. 14074; PIEZO1 mouse antibody, Invitrogen cat. MA5-32876; GAPDH mouse antibody, CST cat. 97116S) were added to the membrane and incubated on a rocker at 4°C overnight. After three washes with TBST, secondary antibody (HRP-Goat pAb to Rabbit IgG, abcam cat. AB205718; HRP-Goat anti-mouse IgG, Invitrogen cat. 62-6520) was added and incubated at room temperature for one hour. Western blot substrate (Bio-Rad cat. 170-5061) was added and membranes were imaged on an Invitrogen iBright imaging system.

### Survival analysis

Survival analysis was performed using the R2 Genomics Analysis and Visualization Platform (Tumor Rhabdomyosarcoma – Williamson dataset; Chromosomal_translocation_negative subgroup; identifier: *ps_avgpres_emtab1202ebi101_u133p2*).

### Calcium influx imaging

For calcium influx imaging, cells were seeded on 8-chamber cover slides to achieve 30-70% confluency on the day of assay. For each chamber, 500 µL culture media containing 2 µL Fluo-4 AM (abcam cat. Ab241082) and 1 µL SPY555-Actin (Cytoskeleton cat. CY-SC202) was added to cultured cells and incubated for 1 hour. If drug treatment was performed at this step, drugs were added and diluted to working concentration in media prior to the addition of dyes. After incubation, staining solution was discarded and cells were washed twice with PBS, and 500 culture media without pheno-red (DMEM containing 10% FBS and 1% penicillin/streptomycin) was added to each well and imaged on a Nikon inverted fluorescent microscopy. Calcium influx intensities were assessed using MATLAB custom-made algorithm.

### Drug Treatment

All small molecule drugs used in this study were purchased from Selleckchem.com. For spheroid invasion assay, 5µM Dooku1 (cat. 2953) and 5µM GsMTx4 (cat. P1205) were added to culture media when the spheroids are transferred onto collagen gel. For calcium influx imaging, 100 nM verteporfin (cat. S1786) and 5mM Y-27632 (cat. S6390) were added and diluted to working concentration in culture media before actin and calcium dye was added.

### AI Disclosure

During the preparation of this manuscript, Gemini (Google) used for proofreading to improve language and readability. After using this tool, the authors reviewed and edited the content as needed and take full responsibility for the scientific integrity and final version of the manuscript.

## Acknowledgement

We thank generous supports from Tower Cancer Research Foundation, Rally Foundation, Infinite Love for Kids Fighting Cancer Foundation, and Alex’s Lemonade Stand Foundation

## Back Matter

### Contribution

Y.P.: Data curation, supervision, data analysis, writing of the manuscript, review, and editing; J.K.: Data curation, writing; B.M.W.: Data curation; E.E.: Data curation; J.P.: Conceptualization, supervision, grant acquisition, data analysis, writing, review, and editing.

### Funding

This research was funded by the Tower Cancer Research Foundation, Rally Foundation, Infinite Love for Kids Fighting Cancer Foundation, and Alex’s Lemonade Stand Foundation.

### Institutional Review Board Statement

Ethical review and approval were waived for this study, as it utilized exclusively de-identified, publicly available clinical datasets (GSE55186, GSE47198, and R2 Genomics Platform) and established commercial cell lines.

### Informed Consent Statement

Not applicable. Patient data used in this study were obtained from public repositories where informed consent was previously obtained by the original investigators.

### Data Availability Statement

The data presented in this study are available in the manuscript and figure legends. Publicly available datasets were analyzed in this study; ChIP-seq data can be found at GSE55186, and RNA-seq data can be found at GSE47198. Survival analysis was conducted using the R2 Genomics Analysis and Visualization Platform (identifier: ps_avgpres_emtab1202ebi101_u133p2)

